# Early-stage lung adenocarcinoma is driven by an injury-associated, plastic cell state dependent on a KRAS-ITGA3-SRC axis

**DOI:** 10.1101/2024.02.27.582165

**Authors:** Aaron L. Moye, Antonella F. M. Dost, Robert Ietswaart, Shreoshi Sengupta, VanNashlee Ya, Chrystal Aluya, Caroline G. Fahey, Sharon M. Louie, Margherita Paschini, Carla F. Kim

## Abstract

Glycine 12 mutations in the GTPase KRAS (KRAS^G12^) are a known initiating event for lung adenocarcinoma (LUAD) with broad clinical relevance. KRAS^G12^ mutations promote cell-intrinsic rewiring of the lung alveolar type II progenitor (AT2) cells, but to what extent such changes interplay with pathways essential for lung homeostasis and cell fate is unclear. We used single-cell RNA-seq (scRNA-seq) from AT2-mesenchyme organoid co-cultures, mouse models, and stage IA LUAD patient samples to identify conserved regulators of AT2 cell transcriptional dynamics and the impact of KRAS^G12D^ with temporal resolution. In AT2^WT^ organoids, a transient injury/plasticity state preceded AT2 self-renewal and AT1 differentiation. Early-stage AT2^KRAS^ cells exhibited perturbed gene expression dynamics most noted by retention of the injury/plasticity state. At later time points in tumorigenesis, AT2^KRAS^ cells consisted of heterogeneous populations that could be defined by either the injury state or high expression of an AT2 cell signature. The injury state in AT2^KRAS^ cells of LUAD in patients, mice, and organoids was distinguishable from AT2^WT^ states by altered receptor expression, including co-expression of ITGA3 and SRC. The combination of clinically relevant KRAS^G12D^ and SRC inhibitors to target the oncogenic injury cell state impaired AT2^KRAS^ organoid growth. Thus, an injury/plasticity signature characterized as an essential step in lung repair is used during alveolar cell self-renewal and during initiation and progression of LUAD. Early-stage lung cancer may be susceptible to intervention by targeting the oncogenic-specific nature of this cell state.

## Introduction

LUAD is the leading cause of cancer-associated death worldwide (*1*). Oncogenic mutations in KRAS are present in 30% of LUAD patients (*2*, *3*). KRAS^G12^ variants are sufficient to initiate LUAD in genetically engineered mouse models (GEMMs) and accurately model human LUAD (*4*). Surfactant-producing AT2 cells have important progenitor functions during injury repair via differentiation into alveolar type I (AT1) cells, the pneumocytes that perform gaseous exchange. Recent studies have defined injury-associated intermediate states in AT2-to-AT1 differentiation (*5*–*7*). We previously showed that KRAS^G12D^ causes transcriptome rewiring in AT2 cells during LUAD development (*8*). Others have also examined the transcriptional (*9*, *10*), epigenetic (*11*), and metabolic (*12*) landscapes of different stages of LUAD. However, there are still significant questions regarding early-stage LUAD: the interplay between pathways used by AT2 cells in homeostasis, repair, and oncogenesis and how cell-intrinsic KRAS-driven changes alter signaling with the microenvironment is largely unknown.

Identifying cellular states at different stages of LUAD using powerful model systems may provide insight into mechanisms of LUAD progression and novel targets for therapeutic intervention. Small molecule inhibitors directly targeting oncogenic KRAS^G12C^, such as adagrasib and sotorasib, MRTX113 targeting KRAS^G12D^, and numerous other pan or variant-specific KRAS-targeting drugs are entering the clinic (*13*, *14*). The discovery of variant-specific KRAS inhibitors is a major breakthrough, yet clinical trial data has revealed that only a subset of indicated KRAS mutant patients respond to treatment (*15*, *16*), highlighting the need to find novel combination treatments to improve efficacy and overcome resistance mechanisms. Significant improvements in early-stage cancer detection using low-dose computed tomography (LDCT) (*17*) and blood-based detection assays (*18*, *19*) will ultimately result in more patients being diagnosed, increasing the demand for early-stage therapeutics. To address these gaps, we sought to identify transcriptional dynamics downstream of activation of oncogenic KRAS expression in AT2 cells.

### Lung AT2 progenitors activate a transient injury/plasticity state prior to both AT2 self-renewal and AT1 differentiation

To identify transcriptional changes specific for KRAS-mutant LUAD initiation at the earliest stages, we generated and analyzed scRNA-seq data from control *Rosa26^YFP^* AT2 organoids (henceforth, AT2^WT^), *Kras^G12D/WT^ p53^flox/flox^ Rosa26^YFP^* (henceforth, AT2^KRAS^) AT2 organoids, and their co-cultured non-cancerous mesenchymal cells. We used our previously described 3D organoid culture combined with Cre-mediated, adenoviral in vitro induction (Fig. 1A) (*8*). AT2 (YFP^POS^) organoids and mesenchymal (YFP^NEG^ EPCAM^NEG^) cells were collected using fluorescence-activated cell sorting (FACS) at 4, 7, 10, and 14 days after infecting cells with Ad5-CMV-Cre (fig. S1A). The scRNA-seq data was generated using the 10X Genomics 3′ platform and preprocessed for further analysis (n=71,252) (fig. S1, B-D). Transcriptionally distinct states were identified using differential expression (DE) analysis on the combined AT2^WT^ and AT2^KRAS^ organoid data binned by genotype and time point (fig. S1D and table 1). After DE genes were filtered for transcription factors (TFs) we observed that cells from each time point had a distinct TF profile, confirming that both AT2^WT^ and AT2^KRAS^ organoids are transcriptionally rewired in a temporally-defined manner (fig. S1E). We previously showed SOX9 and loss of AT2 marker gene expression are hallmarks of transcriptional changes in AT2 cells early after *Kras* mutation (*8*). In agreement, *Sox9* expression was absent in all AT2^WT^ organoid time points, was expressed exclusively in AT2^KRAS^ organoids after Day 7, and correlated with a decreased expression of AT2 lineage-defining genes *Etv5*, *Lyz2,* and *Nkx2.1* (Fig. 1B). We benchmarked our time course data by comparing the expression of genes reported to promote tumorigenesis in *Kras*-mutant GEMMs. In AT2^KRAS^ organoids, *Tigit*, *Cldn4*, and *Itga2*, markers of a high plasticity cell state (HPCS) (*10*), and *Lgr5* were expressed at all AT2^KRAS^ time points (Fig. 1B) (*20*). AT2^KRAS^ organoids also expressed *Krt8*, a marker of transitional injury-associated cell state between AT2 cells and AT1 cells, observed during lung regeneration (*7*). We noticed that many of the HPCS genes are also genes that comprise the injury-associated transitional state and thus termed this an injury/plasticity state. Examining this set of genes, we noted their transient expression in AT2^WT^ organoids, particularly present at Day 4 and absent at later time points when cells with high expression of either AT2 or AT1 cell signature genes were present. In contrast, AT2^KRAS^ organoid cells retained this injury/plasticity state across all time points (Fig. 1B).

**Figure 1.**
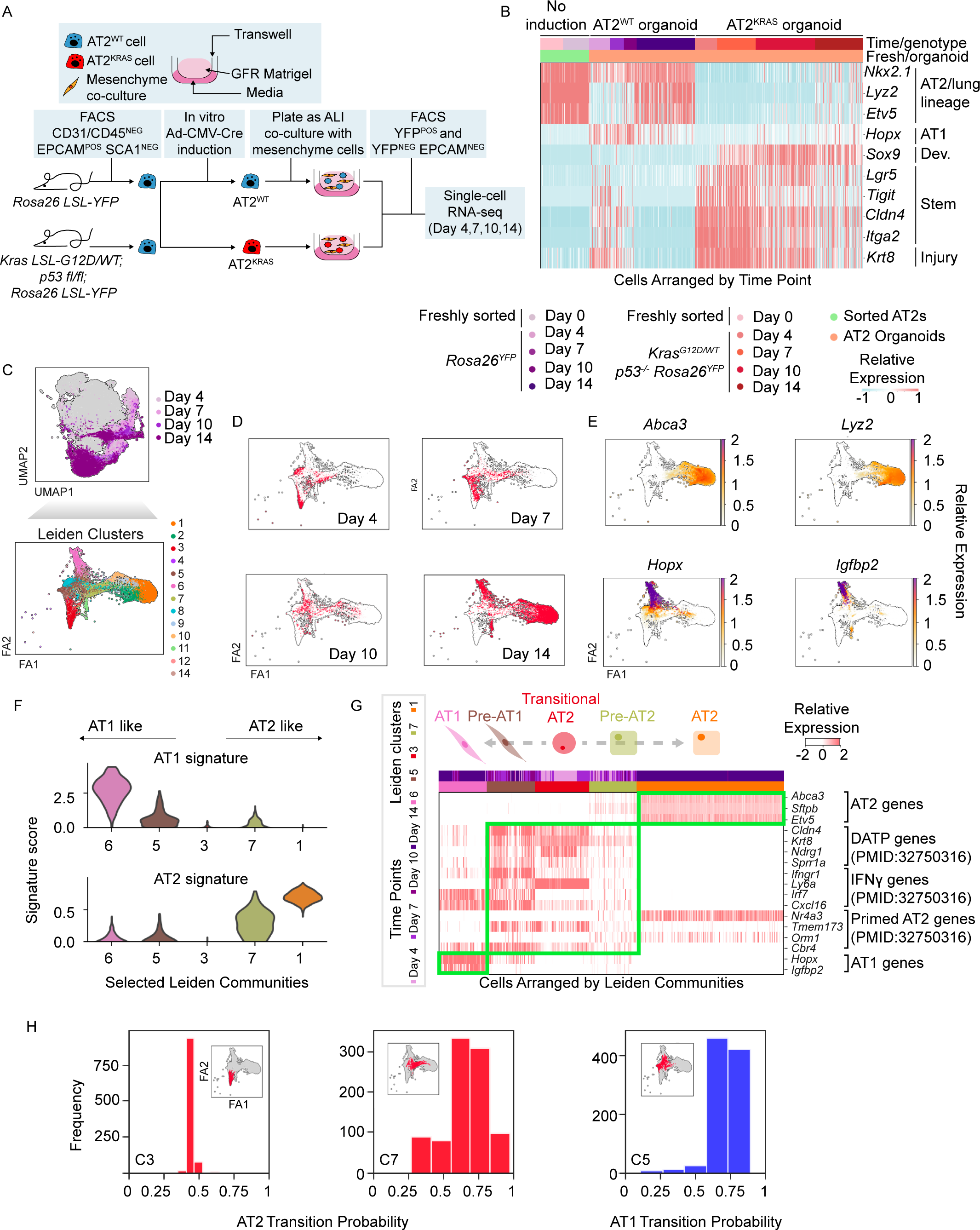
(A) Schematic representation of the AT2-mesenchyme organoid co-culture time course experiment. (B) Heatmap of gene expression in organoid time course data relevant to AT2, AT1, development, stem cell, and injury response genes. “No induction” are primary AT2 cells collected from each mouse genotype (Y and KPY) prior to viral induction or culturing as organoids. (C) FA2 representation of filtered single cells subset from AT2^WT^ organoid data and their corresponding Leiden community. (D) FA2 representation of filtered single cells subset from AT2^WT^ organoid data and their corresponding time point. (E) FA2 representations of filtered single cells subset from AT2^WT^ organoid data and the relative expression of either AT1 or AT2 specific genes. (F) AT2 and AT1 signature score level per community using a violin plot (y-axis, signature score; x-axis, Leiden community). (G) Heatmap of relative gene expression in the AT2^WT^ organoid time course data relevant to AT2, AT1, and injury response gene expression signatures. Cells are arranged with C3 (Day 4 AT2^WT^ organoids) in the middle, left is the AT1 fate, and right is the AT2 fate path. (H) Transition probabilities of cells in selected AT2^WT^ Leiden communities (x-axis; labeled transition probability, y-axis; binned frequency of occurrence). Inset highlights the cell states whose probabilities are being represented within the FA2 space.

To further examine the transcriptional changes of AT2^WT^ cell states, we subset the epithelial component of our AT2^WT^ organoid data and performed Leiden community detection (Fig. 1C and fig. S2A). AT2^WT^ organoids initially formed a transcriptionally homogenous cell state at Day 4 before establishing two distinct terminal states by Day 14. Over the 14 days examined, two distinct AT2^WT^ endpoints formed with either an AT1 cell (Leiden community C6) or AT2 cell signature (C1) with distinct intermediate cell states (C5 and C7) preceding both fates, respectively (Fig. 1, D-F and fig. S2, B and C). Along the AT1 fate (C5 and C6), we observed the expression of *Hopx^POS^ Igfbp2^NEG^* cells and *Hopx^POS^ Igfbp2^POS^*, which are markers for stem-like and terminally-differentiated AT1 cells, respectively (*21*), and cells with AT2 fate had little to no expression of either gene (Fig. 1E). Furthermore, the early-stage Day 4 AT2^WT^ state (C3) had little to no AT1 or AT2 signature score indicating that AT2^WT^ cells enter a lineage plastic state as part of homeostatic regeneration (Fig. 1F). We found elevated *Tead1* and *Wwrt1* (encodes for TAZ) expression in the AT1 fate, consistent with recent observations that YAP/TAZ has a vital role in AT1 cell identity (*22*, *23*) (fig. S2D).

Next, we further investigated the potential relationships between AT2^WT^ organoid states. First, we examined the association between our temporally defined states and recently discovered AT2 injury states, identified as precursors to AT1 differentiation (*5*, *7*). Our analysis revealed that “DATP” (*Cldn4, Krt8, Ndgr1, Sprr1a*), “IFNγ” (*Ifgnr1, Ly6a, Irf7, Cxcl16*), and “Primed AT2” (*Nr4a3, Tmem173, Orm1, Cbr4*) gene signatures were activated in early (C3) and intermediate (C5 and C7) AT2^WT^ states but were suppressed by Day 14 (Fig. 1G). Furthermore, these injury signatures negatively correlated with AT2 signature expression, suggesting that injury signatures are a precursor for AT2 self-renewal and correlates with lineage plasticity. We next tested the fate commitment probabilities of early (C3) and intermediate (C5 and C7) states in AT2^WT^ organoids. We computationally determined the transition probability of C5 and C7 using Palantir, which captures the continuity in cell states modeled as a stochastic process, then assigns a probability of differentiating into pre-defined terminal states (*24*). Start and end points were defined using our temporal information and gene expression, with C3 (Day 4) being the start point and C6 (AT1, Day 14) and C1 (AT2, Day 14) being the endpoints. C3 had approximately a 50% probability of transitioning to the AT1 or AT2 state, supporting our model that a lineage-plastic state is found in AT2^WT^ organoids with AT1 and AT2 differentiation potential. For the intermediate states, the majority of C7 state were predicted to transition to the AT2 state; in contrast, nearly all C5 state were predicted to transition to the AT1 state (Fig. 1H). Hence, within our temporal AT2^WT^ organoid data we identified early-stage activation of a transient, injury/plasticity (AT2-sig^LOW^ AT1-sig^LOW^) state which precedes the formation of fate-committed intermediate states and endpoint AT2 and AT1 states. Lastly, we sought to better understand the TF regulators of fate choice in AT2^WT^ organoids. We subset AT1 and AT2 intermediate states, performed DE analysis, and filtered for TFs (p-value < 0.05 and log_2_ fold change > 1) using a custom-curated reference list (*25*, *26*) (table 2) (fig. S2E). *Nfe213* was amongst the top DE TFs in C5 and was previously identified as DE in E15–E18 pre-ATI cells (*27*), consistent with C5 activating a *Hopx^POS^ Igfbp2^NEG^* AT1 state. In contrast, the AT2-lineage defining TF *Etv5* was DE in C7 (*28*) (fig. S2F). Collectively, our temporal data supports an unappreciated model where AT2^WT^ cells activate a transient, injury/plasticity state defined by AT2-lineage suppression and activation of injury-responsive gene programs, followed by the commitment to an AT2 or AT1 fate outcome via transcriptionally distinct AT2- and AT1-intermediate states. We also observed no evidence of Day 14 AT2 organoids transdifferentiating to the AT1 state based on FA2 representation of the data, arguing that the injury/plasticity state is an important intermediate step in AT2^WT^ self-renewal and AT1 differentiation (Fig. 1E).

### Kras G12D promotes distinct temporal subpopulations defined by AT2-lineage and injury/plasticity signatures

Having identified an injury/plasticity state in AT2^WT^ cells undergoing self-renewal, we wanted to determine if this state is co-opted during tumorigenesis. We analyzed the AT2^KRAS^ subset of the organoid data, re-processed, and identified Leiden communities (Fig. 2A, fig. S3A, and table 3). At Day 4, AT2^KRAS^ organoid cells formed a single community (C4) similar to AT2^WT^ organoids. However, unlike AT2^WT^ cells, AT2^KRAS^ cells had dysregulation in subsequent time points, lacking the AT2 or AT1 fate commitment (Fig. 2B). Notably, AT1 marker genes *Hopx* and *Igfbp2* as well an AT1 signature were suppressed in AT2^KRAS^ cells, and AT2 marker genes (*Abca3*, *Lyz2* and a multiple-gene AT2 signature) were only robustly expressed in a subpopulation of Day 14 organoid cells (Fig. 2, B-D). In contrast to AT2^WT^ cells with transient expression of injury/plasticity genes during organoid culture, these transcriptional signatures were prolonged beyond Day 7 in AT2^KRAS^ organoids (Fig. 2E). At Day 14, AT2^HIGH^ C5 cells re-activated canonical AT2 genes *Abca3*, *Sftpb*, and *Etv5* while suppressing injury/plasticity genes (Fig. 2E). In contrast, Day 14 AT2^LOW^ C2 cells maintained the lineage plastic state characterized by continued expression of injury response genes, suppression of AT2 genes, and elevated expression of *Sox9* (Fig. 2E and fig. S3B). To further examine if injury/plasticity and AT2 lineage signature can be used to temporally define AT2^KRAS^ heterogeneity, we subset mid- and late-stage AT2^KRAS^ populations and calculated signature scores. Indeed, distinct cell states were identifed in AT2^KRAS^ progression based on injury/plasticity and AT2 signature. Furthermore, cells with high injury/plasticity signature had low AT2 signature (Fig. 2F). Lastly, we determined the probabilities of AT2^KRAS^ cell states to form either the AT2 low or high states, where Day 4 C4 state was defined as the starting point. The vast majority of Day 4-10 states, both injury/plasticity^HIGH^ and AT2^HIGH^, were predicted to transition to Day 14 AT2^LOW^ state, suggesting that reactivation of the AT2^HIGH^ state is not an intrinsically favorable outcome in AT2^KRAS^ progression (Fig. 2G); Despite this, reactivation of the AT2^HIGH^ state was a strong feature in the latest time points in our study, suggesting non-cell-intrinsic influence on cell states. Overall, KRAS-mutant AT2 cells co-opt the injury/plasticity state beyond the transient nature observed in AT2^WT^ organoids, resulting in distinct cellular states across early-stage and late-stage time points, delineated by injury/plasticity or AT2-lineage signatures.

**Figure 2.**
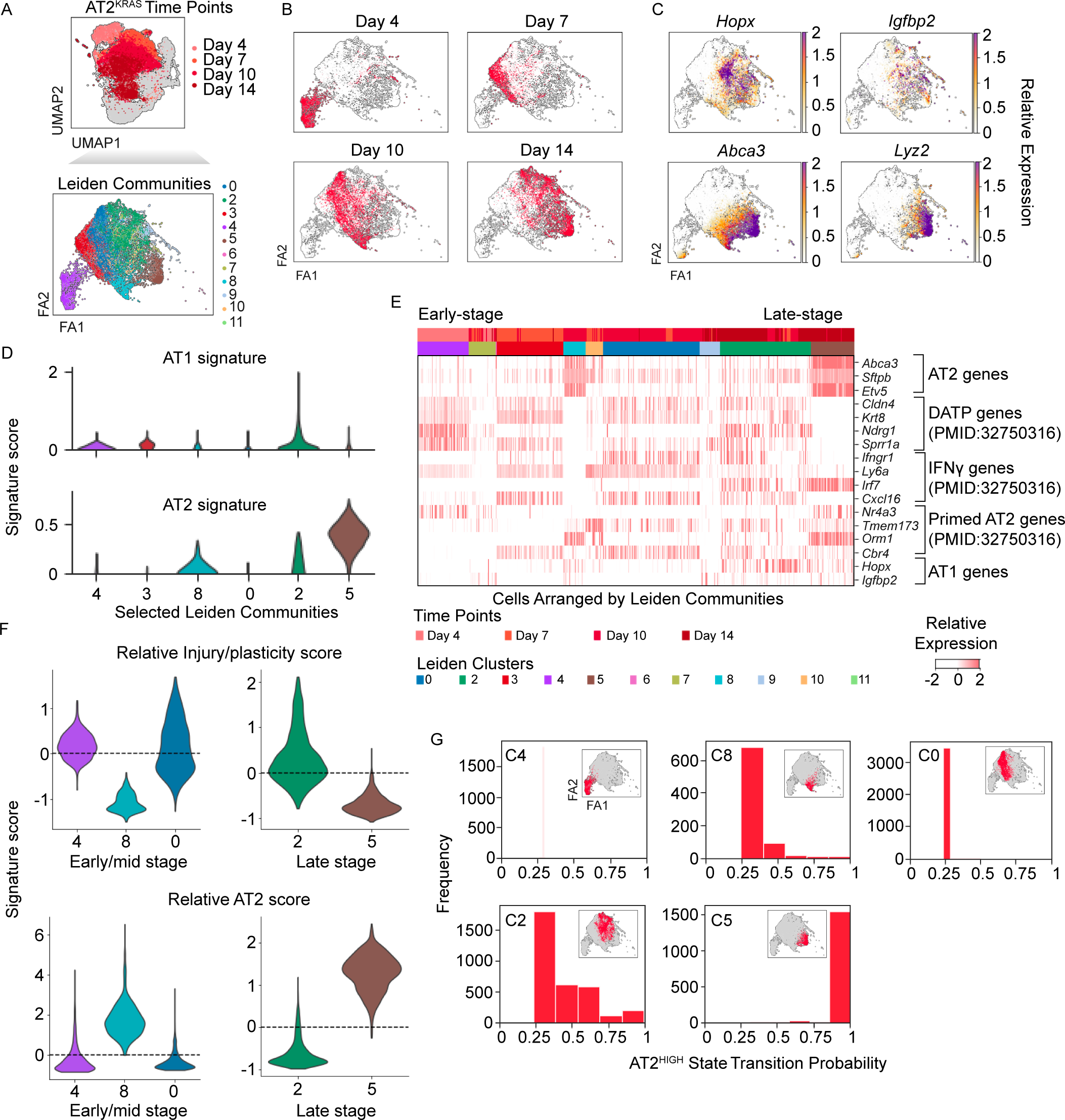
(A) FA2 representation of filtered single cells subset from AT2^KRAS^ organoid data and their corresponding Leiden community. (B) FA2 representation of filtered single cells subset from AT2^KRAS^ organoid data and their corresponding time point. (C) Relative expression of AT1 (*Hopx* and *Igfbp2*) and AT2 (*Abca3* and *Lyz2*) genes within the AT2^KRAS^ FA2 space. (D) AT2 and AT1 signature score level per community using a violin plot (y-axis, signature score; x-axis, Leiden community). (E) Heatmap of relative gene expression in the AT2^KRAS^ organoid time course data relevant to AT2, AT1, and injury response gene expression signatures. Cells are arranged by time point contributions in each Leiden community; early (left), late (right). (F) Relative injury/plasticity and AT2 signature score level per community using a violin plot (y-axis, signature score; x-axis, Leiden community). (G) Transition probabilities of cells in selected AT2^KRAS^ Leiden communities (x-axis; labeled transition probability, y-axis; binned frequency of occurrence). Inset highlights the cell states whose probabilities are being represented within the FA2 space.

### AT2^KRAS^ states are temporally conserved in vivo and in stage IA human LUAD

Having identified an injury/plasticity state in AT2^WT^ self-renewal and in the context of oncogenic *Kras* in orgnoids, we sought to validate our observation in vivo by generating a scRNA-seq time course dataset using the *Kras^G12D/WT^ p53^flox/flox^ Rosa26^YFP^* GEMM. Mice were treated intratracheally with an Ad5-SPC-Cre, resulting in tumor initiation specifically in AT2 progenitors (Fig. 3A). AT2 cells were collected using FACS at 4, 7, 10, and 16 weeks post-infection, covering the transition from hyperplasia to adenocarcinoma (*4*) (fig S4A). Furthermore, mice were induced so that AT2 cells could be collected and scRNA-seq libraries generated at the same time (materials and methods), allowing for direct comparison of time points without the need for batch correction. The scRNA-seq libraries were generated using the 10X Genomics platform, and the data processed for downstream analysis (n=3,125 cells) (Fig. 3B and fig. S4B). Consistent with our observations in organoids, AT2 lineage gene expression negatively correlated with development and injury response genes (fig. S4C). To further examine temporal AT2^KRAS^ heterogeneity, Leiden communities were identified (Fig. 3B and fig. S4D). Consistent with our organoid data, at the late-stage week 16 time point we observed the formation of distinct AT2^HIGH^ (C7) and AT2^LOW^ (C3) states (Fig. 3C). AT2^LOW^ cells had elevated injury-associated gene expression while AT2^HIGH^ cells suppressed injury-associated gene expression (Fig. 3C). Next, we wanted to determine if injury/plasticity and AT2 signatures can define distinct mid- and late-stage cell states in vivo. In agreement with our organoid data, injury/plasticity^HIGH^ AT2^LOW^ and injury/plasticity^LOW^ AT2^HIGH^ populations were observed at mid and late-stage time points (Fig 3D) confirming a conserved interplay between injury/plasticity and AT2-lineage gene expression dynamics in LUAD progression. Lastly, to establish human relevance we reanalyzed our scRNA-seq data from two stage IA, Kras-mutant LUAD patients (*8*). Preprocessing the data, we subset human AT2^WT^ and AT2^KRAS^ cells along with previously uncharacterized mesenchymal cells (COL1A1^POS^ VIM^POS^) (n=3,331 cells) (Fig. 3E and fig S4, E-G). In agreement with our organoid and in vivo data, early-stage human AT2^KRAS^ cells have reduced AT2 lineage-defining gene expression. Furthermore, injury/plasticity gene expression signatures, including *CLDN4* and *KRT8* were robustly upregulated in human AT2^KRAS^ cells (Fig. 3F and fig. S4G). This data demonstrates that AT2^KRAS^ cells utilize a normally transient injury/plasticity^HIGH^ AT2^LOW^ state for AT2 self-renewal, which promotes LUAD progression and heterogeneity, and is conserved in AT2 cells in vivo, in organoids, and human patients with early-stage LUAD (Fig. 3G).

**Figure 3.**
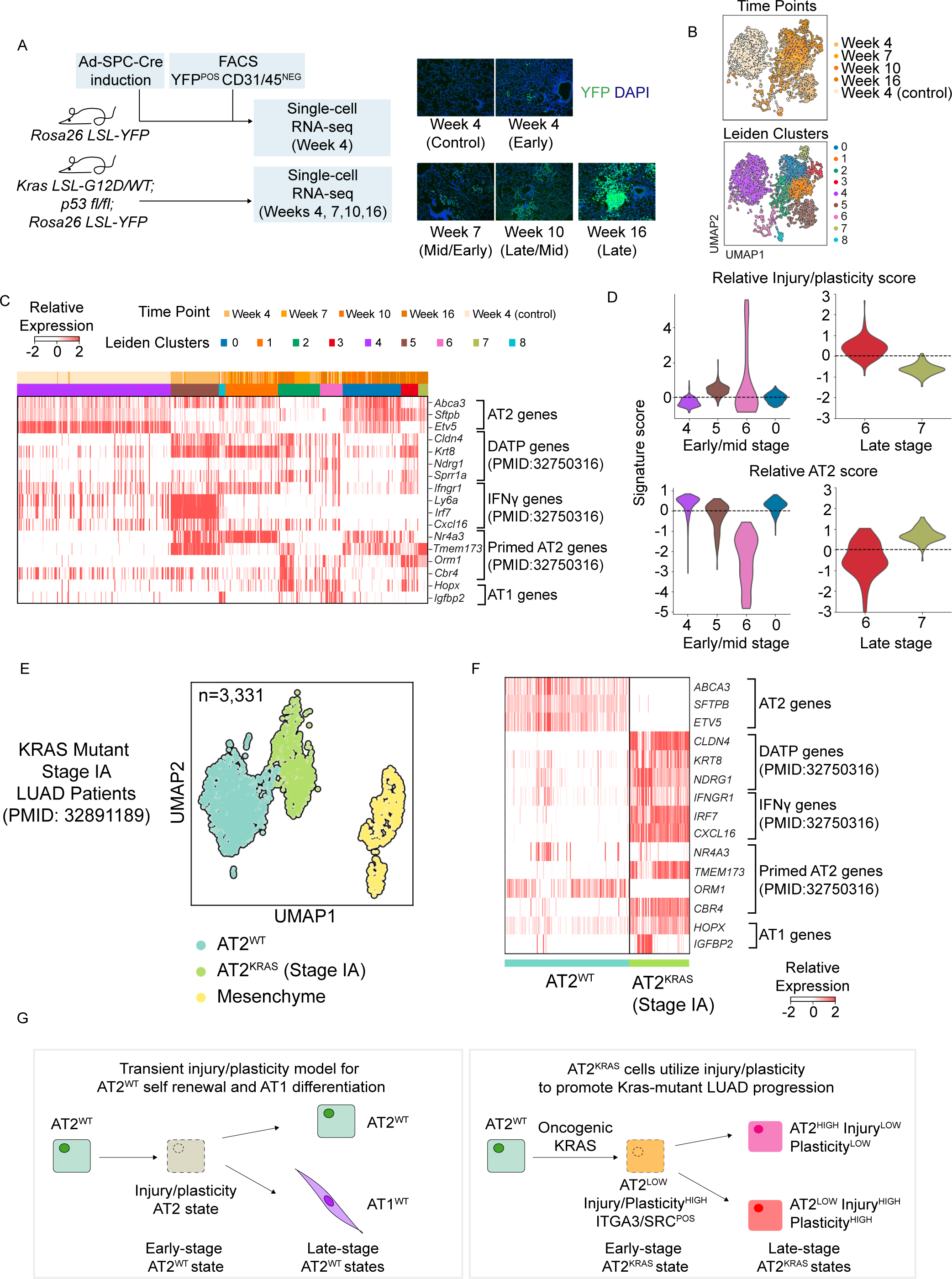
(A) Schematic representation of the in vivo time course experiment. (B) UMAP representation of filtered single cells from the in vivo experiment, their corresponding time point, and Leiden communities. (C) Heatmap of relative gene expression in the in vivo time course data relevant to AT2, AT1, and injury response gene expression signatures. Cells are arranged by time point contributions in each Leiden community; early (left), late (right). (D) Relative injury/plasticity and AT2 signature score level per community using a violin plot (y-axis, signature score; x-axis, Leiden community). (E) UMAP representation of filtered single cells from previously published stage IA LUAD patients data (*8*) and their corresponding Leiden community (this paper). (F) Heatmap of relative gene expression in the human stage IA data relevant to AT2, AT1, and injury response gene expression signatures. Cells are arranged by Leiden community. (G) Proposed model for AT2^WT^ and AT2^KRAS^ transcriptional cell states over time conserved in murine organoids and in vivo. A conserved injury/plasticity state precedes both AT2 self-renewal and AT1 differentiation, which is co-opted by oncogenic KRAS to drive LUAD initiation, heterogeneity, and progression. Importantly, the cell states have distinct transcriptomes in each time point and condition (represented by different colors).

### A KRAS-ITGA3-SRC axis drives the earliest stages of LUAD initiation

Having determined that an injury/plasticity state is shared by self-renewing and tumorigenic AT2 cells, we next sought to determine if *Kras*-driven cells utilize pathways to drive tumorigenesis that are distinct from normal progenitor cell activity and differentiation. We performed DE analysis based on Leiden communities in our AT2 organoid, GEMM, and stage IA human data (Fig. 4A, fig. S5A, and table 4). DE genes were collected from early-stage states from each dataset: Week 4 GEMM in vivo, Day 4 organoids, and AT2^KRAS^ cells from stage IA LUAD patients. Lastly, we filtered for statistically significant (p-value < 0.05) DE genes identified receptors (*29*) (Fig. 4A). Our analysis unbiasedly identified eight receptors differentially expressed by early-stage Kras-mutant cells across all three model systems: *OCLN, ITGA3, ADIPOR1, PLXNB2, CLDN4, ST14, ITGB1,* and *LSR* (Fig. 4B). Importantly, in the GEMM and organoid datasets with temporal resolution, the majority of identified receptors had the highest relative expression in early-stage time points, and elevated relative expression in AT2^KRAS^ cells compared to AT2^WT^ cells in our stage IA human data (Fig. 4B). In a separate analysis we used Genewalk, a machine-learning (ML) approach to find highly relevant genes and Gene Ontology (GO) terms in early-stage AT2^KRAS^ cells. The graph-based ML generates a biomedical knowledge graph to identify context-specific interconnections between input genes and biological pathways (*30*). We used DE genes upregulated in the early-stage AT2^KRAS^ (C8) relative to the AT2^WT^ (C10) communities as input for ML analysis (table 5). Genewalk identified integrin subunits *Itga3* and *Itgb1* as receptors with the highest gene-gene and gene-GO interconnections (Fig. 4C) within our identified receptors, prompting us to further investigate the conserved role of integrins in early-stage AT2^KRAS^ cells. Integrins are heterodimeric receptor complexes that comprise an alpha and beta subunit. There are 24 known human integrin heterodimers that have evolved specialized functions (*31*–*33*). We chose to focus on the alpha subunit *Itga3* because the role of α3 integrins in early-stage LUAD is poorly understood. To validate the observation that early-stage AT2^KRAS^ cells have high expression of *Itga3*, RNAScope (*34*), an in situ hybridization technique, was performed using Itga3 specific probes combined with IF staining for YFP, a marker of AT2^KRAS^ cells, and the AT2 marker SPC. Consistent with scRNA-seq data, *Itga3* had the strongest signal in YFP^POS^ cells at 4 weeks (early) compared to 10 weeks (mid-stage) time point in vivo, with little to no *Itga3* signal in wildtype AT2 cells (YFP^NEG^ SPC^POS^) (Fig. 4D). After confirming that high *Itga3* expression is specific to early-stage AT2^KRAS^ cells, we examined gene-GO associations identified by our ML analysis. The top GO hit for *Itga3* was “integrin alpha3 beta1 complex” (Fig. 4E). Intriguingly, *Itgb1* was also a conserved receptor hit (Fig. 4B), suggesting a possible function of α3β1 integrin in early-stage AT2^KRAS^ cells. We observed a positive correlation between the expression of *Itga3* and the injury/plasticity signature at early-stage AT2^KRAS^ time points, when *Itga3* levels were most significantly DE, and also at mid and later stages (fig. S5B).

**Figure 4.**
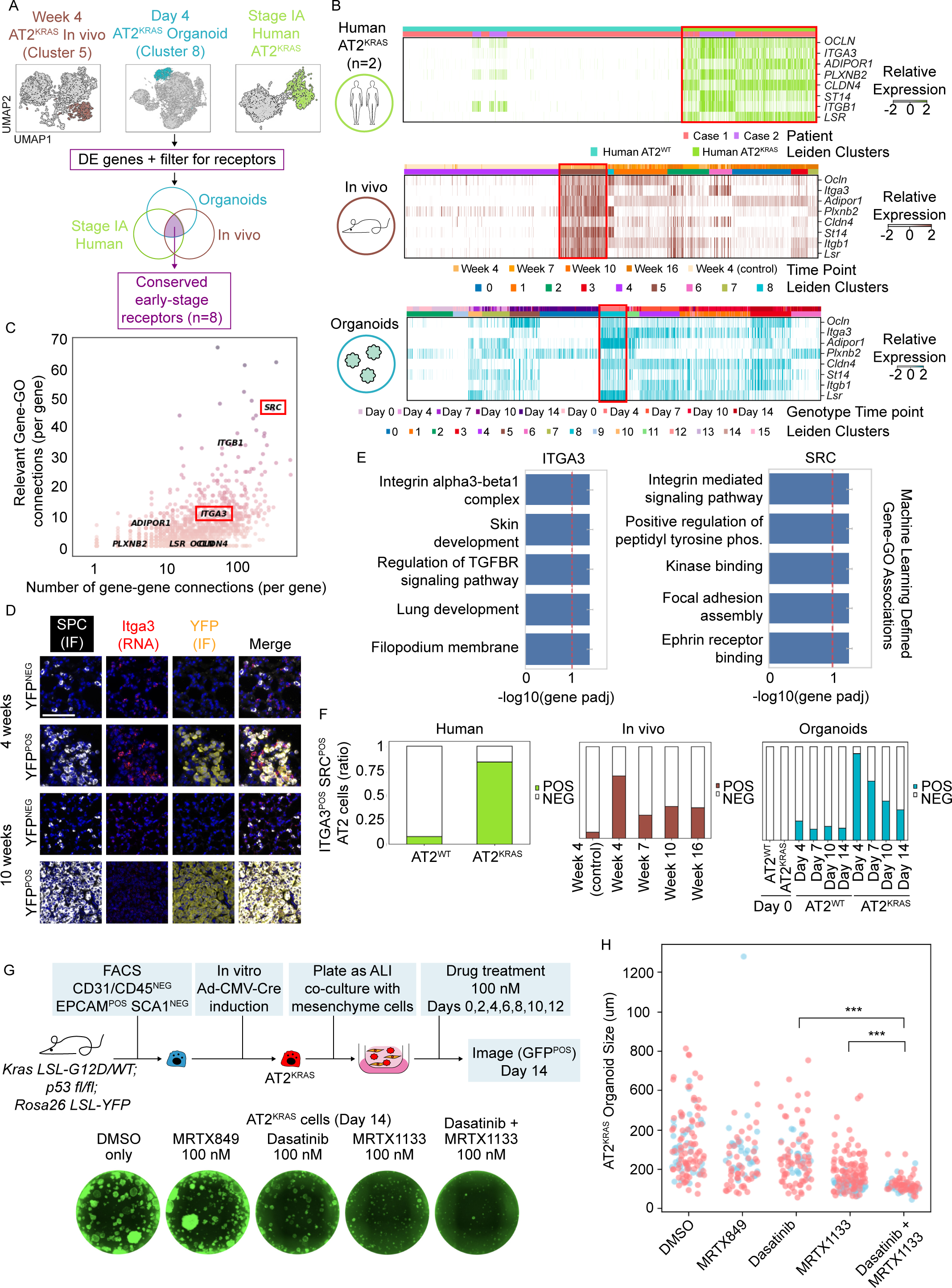
(A) Analysis design to identify early-stage AT2^KRAS^ receptors conserved across species and model system examined in this study. (B) Relative expression of identified receptors in the AT2 organoids, in vivo, and human patient data relevant, using a heatmap (x-axis; Leiden communities arranged by time point contributions, y-axis; conserved receptors). (C) Interconnections between DE genes in early-stage AT2^KRAS^ organoids identified using machine learning, represented as a dot plot (x-axis; number of gene-gene connections, y-axis; number of gene-GO term connections). Genes selected for further study are highlighted with red box. (D) RNAScope analysis in 4-week and 10-week murine lung section following in vivo induction with Ad5SPC-Cre. Scale bar = 100 um. (E) Bar plot of context-relevant GO terms associated with *Itga3* and *Src* in early-stage AT2^KRAS^ organoids, identified using machine learning. GO terms are arranged by statistical significance. (F) Bar plot representing the percentage of Itga3^POS^ Src^POS^ cells in each Leiden community in the AT2 organoid, in vivo, and human patient datasets. (G) Experimental design of AT2^KRAS^ organoid co-cultures treated with DMSO (vehicle), MRTX849, MRTX1133, Dasatinib, or MRTX1133/Dasatinib combination, and representative images of D14 AT2^KRAS^ organoids from each treatment group. (H) Quantitation of Day 14 AT2^KRAS^ organoid size (diameter, um) in each treatment group; A total of 80-137 organoids were measured from two independent experiments, represented as blue and red circles in the graph. ** p < 0.001. DMSO vs. all conditions were statistically significant p < 0.05.

We next probed what may be downstream of integrin signaling in early AT2^KRAS^ cells. Further analysis of our ML results revealed proto-oncogene tyrosine kinase SRC as the fifth most interconnected hit in early AT2^KRAS^ cells (Fig. 4C and table 5). SRC is a known downstream signal of integrins in leukocytes (*31*, *35*), but with uncharacterized function in AT2 cells. The top SRC GO hit identified by our ML analysis was “integrin-mediated signaling pathway” (Fig. 4E). The known connection between SRC and integrin signaling in non-lung cells, the strength of the SRC hit in our ML analysis, and the ML-identified association between SRC and integrin signal made it an ideal candidate for further investigation. We found a positive correlation between *Itga3* and *Src* in AT2 cells at different time points (fig. S5C). Next, we calculated the percentage of AT2 cells double positive for *ITGA3* and *SRC* (relative expression > 0). *Itga3^POS^ Src^POS^* AT2 cells were enriched in early-stage AT2^KRAS^ time points across all models examined in this study (Fig. 4F).

Lastly, we investigated if the KRAS-ITGA3-SRC axis in early AT2^KRAS^ cells could be exploited therapeutically in early-stage LUAD. Small molecule inhibitors of the GDP KRAS-OFF state are emerging, such as sotorasib (LUMAKRAS) and adagrasib (KRAZATI) for KRAS^G12C^ mutant cancers, and MRTX1133 for KRAS^G12D^ mutant cancers (*37*). Based on our findings, we reasoned that combining KRAS^G12D^ and inhibitors of Itga3 or SRC will improve efficacy of KRAS inhibitors. Due to the lack of drugs that directly target ITGA3, we inhibited Src using the FDA-approved small molecule dasatinib (SPRYCEL). AT2^KRAS^ organoids treated at Day 0 with vehicle (DMSO) or adagrasib had little impact on growth because the organoids tested are KRAS^G12D^ mutant. Some impact on the size of AT2^KRAS^ organoids was observed using 100 nM of dasatinib (*38*) or MRTX113 as a monotherapy. In contrast, a combination of dasatinib with MRTX1133 dramatically suppressed AT2^KRAS^ organoid growth (Fig. 4, G and H). Thus, both KRAS^G12D^ and SRC-signaling are important for early AT2^KRAS^ growth and could provide a rationale for combination adagrasib/dasatinib treatment as a potential adjuvant or neo-adjuvant treatment for early-stage KRAS^G12D^ mutant LUAD patients (Fig. 4H).

## Discussion

These data define a conserved injury-associated, lineage-plastic state activated by AT2^WT^ cells prior to AT2 self-renewal that is co-opted by oncogenic KRAS to drive tumorigenesis. It has long been established that AT2 cells have facultative regenerative capacity and AT1 differentiation potential. A model was proposed for lung development in which AT2 and AT1 cells form independently via a bipotent progenitor cell (*39*, *40*). More recently, KRT8^POS^ DATP/PATS, injury-associated cell states have been observed as intermediate states in the process of AT2-to-AT1 differentiation (*5*–*7*). Our data suggest that AT2^WT^ cells do not directly self-renew, but rather undergo transient lineage suppression prior to AT2 self-renewal. Additionally, aberrant retention of this injury/plasticity state defines KRAS G12D AT2 cells. However, we do not observe an AT2^HIGH^ Injury/Plasticity^LOW^ state in our early-stage LUAD patient data, suggesting there are subtle differences in LUAD progression in humans and mice. The AT2^KRAS^ cell-intrinsic injury/plasticity state promotes aberrant cell-cell signaling with the lung mesenchyme possibly via a KRAS-ITGA3-SRC axis. Recent studies have highlighted cell-intrinsic lineage plasticity and injury-associated epigenetic rewiring programs in early-stage pancreatic cancer (*41*, *42*). The role of signals from the non-cancerous mesenchyme in early-stage oncogenesis, however, is poorly understood. The KRAS-mutant injury state is enriched for ITGA3^POS^ SRC^POS^ cells at the earliest stages of LUAD initiation and this state can be targeted with clinically relevant small molecules in organoid co-culture systems, implicating “outside-in” epithelial-mesenchymal integrin signaling and downstream SRC signaling in early LUAD. β3 Integrins have been linked with KRAS to promote fitness and survival of established pancreatic, lung, and breast carcinomas, and can promote resistance to EGFR inhibitors (*43*). Effective inhibitors of integrin signaling remain limited (*32*); therefore, other components of integrin signaling pathways may provide novel therapeutic opportunities. The use of SRC inhibitors like dasatinib as a monotherapy for late-stage NSCLC failed Phase II clinical trials (*44*). Our work provides a rationale for re-testing SRC inhibitors focused on early-stage LUAD in combination with KRAS^G12D^ inhibitors such as MRTX1133. Recent findings showed that inhibiting KRAS induced an AT1-like state in wildtype AT2 cells and LUAD and suggested that alveolar differentiation is a resistance mechanism for therapies targeting the RTK–KRAS axis (*45*). Our results demonstrate that dual RTK-KRAS inhibition potently inhibits early-stage LUAD. We are currently investigating whether the mechanism of the effect of the dual RTK-KRAS inhibition on KP organoid growth is via inducing an AT1 state or reduction of the injury/plasticity state. We cannot rule out the possibility that SRC inhibition might enhance the effect of MRTX1133 by enhancing the ability of MRTX1133 to bind to mutant KRAS. Beyond currently available strategies, this study suggests that targeting the unique injury-associated cell states and the related mesenchymal interactions driven by oncogenic mutations can provide a new means to intervene in early-stage LUAD.

## Acknowledgments

We thank members of the Kim Lab for helpful discussion and feedback. We thank Dr. Ronald Mathieu, Mahnaz Paktinat, and the flow cytometry core facility at Boston Children’s Hospital (BCH), the single-cell core facility at Harvard Medical School (HMS), the Zon lab for the use of their 10X Genomics Chromium Controller, the HMS biopolymers facility, the DFHCC rodent histopathology facility, and the BCH Pathology Department histology lab. This work was supported in part by the Damon Runyon Cancer Research Foundation Postdoctoral Fellowship (no. DRG:2368-19) and a Burroughs Wellcome Fund Postdoctoral Enrichment Program Award (no. 1019903) (A.L.M), the Hope Funds for Cancer Research Postdoctoral Fellowship (S.M.L.), EMBO fellowship ALTF 2016-422 (R.I), R01 HL090136, R01 HL132266, R01 HL125821, U01 HL100402, RFA-HL-09-004, R35HL150876, the Cystic Fibrosis Foundation Award KIM19P0, LONGFONDS Accelerate, project BREATH, Gilda and Alfred Slifka, Gail and Adam Slifka, the Cystic Fibrosis/Multiple Sclerosis Fund Foundation Inc., and the Harvard Stem Cell Institute (C.F.K.).

## Data availability

All raw and processed scRNA-seq data were deposited to the NCBI Gene Expression Omnibus (GEO) and Sequencing Read Archive (SRA) under the accession code GSE253461. Jupyter notebooks containing all bioinformatics analysis are available on GitHub https://github.com/alm8517/Kras_timecourse_study

GEO data will be made publicly available upon publication.

## Methods

### Mouse cohorts

*Kras LSL-G12D/WT; p53 flox/flox* (*46*) mice were crossed to mice homozygous for the Rosa26 LSL-YFP allele to obtain *Kras LSL-G12D/WT; p53 flox/flox; Rosa26 LSL-YFP* mice. Mice were maintained in virus-free conditions. All mouse experiments were approved by the BCH Animal Care and Use Committee, accredited by AAALAC, and were performed in accordance with relevant institutional and national guidelines and regulations.

### Stage IA LUAD patient information

As described previously (*8*), samples of two patients with a stage IA LUAD diagnosis were analyzed in these studies. One patient was female, 74 years old, with a KRAS G12F mutation. The other patient was female, 77 years old, with a KRAS G12V mutation. All patients provided written informed consent. The studies were approved by the UCLA institutional review board.

### In vivo adenovirus infection

Young adult mice (less than 6 months) were infected with 6×10^8^ PFU adenovirus containing the AT2-specific SPC promoter by intratracheal instillation as described previously (DuPage et al., 2009). In this study, only female mice were used. To avoid technical and batch variation when generating scRNA-seq libraries, the mice were induced at different times, allowing cells to be collected via FACS at the same time. The 16-week time point was induced first, followed by inducing the 10-week group six weeks later, the 7-week group three weeks later, and finally the 4-week group three weeks later.

### Generating a single-cell suspension from murine lungs

Mice were anesthetized with avertin, perfused with 10 mL PBS through the trachea, followed by intratracheal instillation of 2 mL dispase (Corning). Lungs were the placed on iced, minced and incubated in 0.0025% DNase (Sigma Aldrich) and 100 mg/ml collagenase/dispase (Roche) in PBS for 45 min at 37 C, filtered through 100 mm and 40 mm cell strainers (Fisher Scientific), and centrifuged at 1000 rpm, 5 min at 4 C. Cells were resuspended in red blood cell lysis buffer (0.15 M NH4Cl, 10mM KHCO3, 0.1 mM EDTA) for 1.5 min, washed with advanced DMEM (GIBCO), and resuspended in PBS/10% FBS (PF10) at 1 million/100 µl.

### Isolation of primary AT2 cells from murine lungs using FACS

Lung single-cell suspensions, see “Generating a single-cell suspension from murine lungs” were incubated for 10 min on ice with DAPI as a viability dye and the following antibodies: anti-CD31 APC, anti-CD45 APC, anti-Ly-6A/E (SCA1) APC/ Cy7 (all Fisher Scientific), anti-CD326 (EPCAM) PE/Cy7 (Biolegend) (all 1:100). Single stain controls and fluorophore minus one (FMO) controls were included for each experiment. FACS was performed using a FACSAria II to collect CD31^NEG^ CD45^NEG^ EPCAM^POS^ SCA1^NEG^ cells. Cells were collected in PBS/10% FBS, and analysis was performed with FlowJo.

### In vitro infection of primary lung epithelial cells using adenovirus and organoid co-culture

Murine AT2 cells (CD31^NEG^ CD45^NEG^ EPCAM^POS^ SCA1^NEG^) cells isolated by FACS as described in ‘‘Isolation of primary AT2 cells from murine lungs using FACS’’ were split into 2 or 3 equal aliquots, or not split, depending on the experiment, pelleted by pulse spin and resuspended in 100 µL MTEC/Plus media (Zhang et al., 2017a) containing 6×10^7^ PFU/ml of Ad5CMV-Cre in 100 µL per 100,000 cells. The cells were incubated for 1 h at 37 C, 5% CO2 in 1.5 mL tubes. Cells were then pelleted by pulse spin and resuspended in 1x phosphate-buffered saline (PBS). This step was repeated twice for a total of three washing steps. Cells were resuspended in DMEM/F12 (Invitrogen) supplemented with 10% FBS, penicillin/streptomycin, 1 mM HEPES, and insulin/transferrin/selenium (Corning) (3D media) at a concentration of 5,000 live cells (trypan blue negative) per 50 µl. As supporting cells, a mix of neonatal stromal/mesenchyme cells was isolated as described previously (Lee et al., 2014). The stromal cells were pelleted and resuspended in growth factor reduced (GFR) Matrigel at a concentration of 50,000 cells per 50 µl. Equal volumes of cells in 3D media and supporting mesenchyme cells in GFR Matrigel were combined and 100 µL were pipetted into a Transwell (Corning). Plates were incubated for 20 min at 37 C, 5% CO2 until Matrigel solidified. Finally, 500 µL of 3D media was added to the bottom of the transwell, and 3D media was changed every two to three days.

### AT2^KRAS^ organoid drug treatment

In vitro induced organoids we plated as described in “In vitro infection of primary lung epithelial cells using adenovirus and organoid co-culture”. Starting Day 0, organoids were grown in 3D media supplemented with 100 nM of dasatinib (SRC inhibitor), MRTX1133 (KRAS G12D inhibitor), MRTX849 (KRAS G12C inhibitor), a combination of dasatinib and MRTX1133, or DMSO only (vehicle). 500 µL of media was added to the base of the transwell. Drug-treated 3D media was changed every two days. Fresh stocks of drug-treated 3D media were made every four days.

### Collecting cells for scRNA-seq using FACS

For in vivo experiments, cells were collected from mice infected with Ad5SPC-Cre at the same time, see “In vivo adenovirus infection”. Single-cell suspensions were generated, see “Generating a single-cell suspension from murine lungs” then incubated for 15 min on ice and protected from light exposure, with 4′,6-diamidino-2-phenylindole (DAPI) as a viability dye and the following antibodies: anti-CD31 PE (BD Biosciences), anti-CD45 APC (Fisher Scientific), anti-CD326 (EPCAM) PE-Cy7 (Biolegend) (all 1:100). Single stain controls and fluorophore minus one (FMO) controls were included for each experiment. FACS was performed on a FACSAria II, cells collected in PBS/10% FBS, and analysis was done with FlowJo.

For organoid experiments. single cell suspensions were obtained at days 4, 7, 10, and 14 for FACS, cells were incubated with EPCAM-PeCy7 (BioLegend) and DAPI (Sigma-Aldrich) for 10 min on ice. A DAPI only control served as the fluorophore minus one (FMO) control for EPCAM. FACS was performed on a FACSAria II and analysis was done with FlowJo.

### Organoid and in vivo scRNA-seq

ScRNA-Seq was performed using the 10X Genomics platform (10X Genomics, Pleasanton, CA). FACS sorted cells from either mice or organoid cultures were encapsulated with a 10X Genomics Chromium Controller Instrument using the Chromium Single Cell A Chip Kit. Encapsulation, reverse transcription, cDNA amplification, and library preparation reagents are from the Chromium Single Cell Library & Gel Bead Kit v3. Briefly, single cells were resuspended in PF10 at a concentration of 1000 cells ml. The protocol was performed as per 10X Genomics protocols without modification (chromium single cell 3 reagent kits user guide v3 chemistry). Total cDNA and cDNA quality following amplification and clean-up was determined using a QubitTM dsDNA HS assay kit and the Agilent TapeStation High Sensitivity D5000 ScreenTape System. Library quality pre-sequencing was determined using Agilent TapeStation and QPCR prior to sequencing. TapeStation analysis and library QPCR was performed by the Biopolymers Facility at Harvard Medical School. Libraries were sequenced using an Illumina NextSeq500 using paired-end sequencing with single indexing (Read 1 = 28 cycles, Index (i7) = 8 cycles, and Read 2 = 91 cycles). Reads were aligned to the mm10-1.2.0 reference genome and count matrices were generated using CellRanger 3.1.0 (10X Genomics).

### Single-cell RNA-seq data analysis

Jupyter notebooks containing all computational analysis in this study can be found on GitHub (https://github.com/alm8517/KRASmutant_TimeSeries). In brief, count matrices generated by CellRanger were read into the Python single-cell analysis environment Scanpy (Wolf et al., 2018). For the organoid co-culture and human data, cells with > 20% mitochondrial content, which correlated with low read count were removed. The data was normalized, logarithmized, and the significant number of principle components determined using in-built Scanpy functions. Data was denoised using Markov Affinity-based, Graph Imputation (MAGIC) using the following settings (Gene to return = all, k = 5, t = 5, n_pca = 30) (van Dijk et al., 2018) followed by nearest neighbor calculation and UMAP dimensionality reduction. Organoid data was subset into epithelial cells and mesenchymal cells for further analysis. For in vivo data, cells with > 10% mitochondrial content, which correlated with low read count, were removed. The data was normalized, logarithmized, and the significant number of principle components was determined using in-built Scanpy functions. Next, we performed nearest neighbor calculations and UMAP dimensionality reduction. Cell types were annotated and epithelial cells were subset for further analysis. A reference list of murine transcription factors was curated from two publicly available datasets; The Animal Transcription Factor Database (Hu et al., 2019) and the TcoF-DB v2 (*26*). Receptor-ligand relationships were curated from the CellPhoneDB database (*47*) and a previously published curated list of receptor-ligand interactions (*29*). AT1 and AT2 marker genes are from the Panglao database (*48*). Relative (early/mid and late) injury/plasticity (*Cldn4, Krt8, Nr4a3, Ifngr1, Ndrg1, Sox9, Hmga2, Itga2*) and AT2 signatures (*Etv5, Lyz2, Abca3*) were calculated using a subset of signature marker genes defined in the figures and associated Jupyter notebooks. Mesenchyme population markers are from various published papers (*49*–*51*). Lung mesenchyme (*COL1A1^POS^*) was subset from the human scRNA-seq data, re-clustered to identify additional heterogeneity, followed by unbiased community detection using the Leiden algorithm. We used mesenchyme marker genes for alveolar fibroblasts (*COL1A1^POS^ PDGFRA^POS^ LGR5^POS^*), pericytes (*PDGFRB^POS^ CSPG4^POS^*), and airway fibroblasts (*LGR6^POS^ ACTA2^POS^*) (*49*, *50*). Machine learning analysis was performed as previously described, using Genewalk v1.5.3 (*30*).

### Staining and IF of organoid cultures and in vivo lung sections

To image whole wells, multiple overlapping images of live organoid cultures were taken at various time points and stitched together using EvosTM FL Auto2 software. Images were processed using FIJI (*53*). To image YFP-positive cells in murine lung section, a lobe of lung was clamped and removed during the preparation of the Ad5SPCCre treated lungs for scRNA-seq.

### RNAScope of in vivo induced murine lung sections

Sectioned lung tissues underwent deparaffinization by incubation with xylene and rehydration in 100% ethanol. The deparaffinized slides were blocked in RNAscope hydrogen peroxide for 10 minutes at room temperature. Fresh 1X Target Retrieval Reagents were prepared by adding 630mL of distilled water to 70mL of 10X Target Retrieval Reagents in a beaker. The Target Retrieval buffer was heated to a mild boil (98-102℃) and maintained at a temperature within the range of 98-102℃ for Target Retrieval. The slides were submerged in the heated 1X RNAscope Target Retrieval solution and boiled for 15 minutes. After Target Retrieval, the slides were washed in distilled water and 100% ethanol before drying at room temperature for 5 minutes. Hydrophobic barriers were created around each section using the ImmEdge hydrophobic barrier pen which were allowed to dry for 5 minutes. The slides were treated with RNAscope Protease Plus and incubated at 40℃ for 30 minutes in the HybEZ Oven. Meanwhile, the 120mL of RNAscope 50X Wash buffer was warmed to 40℃ for 10-20 minutes before being mixed with 5.88L distilled water in a large carboy to prepare 6L of 1X Wash Buffer. The ready-to-use 1X C1 probe (for labeling ITGA3) was warmed for 10 minutes at 40℃ then cooled to room temperature before use. After the RNAscope Protease Plus incubation period, the protease plus was removed from each section and the C1 probe was immediately applied to entirely cover each section. The C1 probe hybridized for 2 hours at 40℃ in the HybEZ Oven. After probe hybridization, the slides were washed in 1X Wash Buffer and stored overnight in 5X Saline Sodium Citrate (SSC) at room temperature. Reagents AMP1, AMP2, AMP3, HRP-C1, and HRP-blockers (RNAscope Multiplex Fluorescent Reagent Kit v2) were equilibrated at room temperature before use. Excess liquid was removed from the slides and each slide received enough drops of RNAscope Multiplex FL v2 Amp 1 to entirely cover the sections. Slides were incubated in the HybEZ Oven for 30 minutes at 40℃ then washed in 1X Wash Buffer before the addition of RNAscope Multiplex FL v2 Amp 2 to each section. The slides were once more incubated in the HybEZ Oven for 30 minutes at 40℃. After a brief rinse in 1X Wash buffer, the slides received enough drops of RNAscope Multiplex FL v2 Amp 3 to entirely cover each section before being incubated in the HybEZ Oven for 15 minutes at 40℃. Meanwhile, the TSA Vivid 570 fluorophore for labeling the C1 probe was prepared to a dilution of 1:1500 in TSA buffer. To develop the HRP-C1 signal, drops of RNAscope Multiplex FL v2 HRP-C1 were added to entirely cover each slide and incubated in the HybEZ Oven for 15 minutes at 40℃. After a brief rinse in 1X Wash Buffer, each section received approximately 150-200 μL diluted fluorophore and the slides incubated for 30 minutes at 40℃. A brief rinse of the slides in 1X Wash Buffer followed incubation and RNAscope Multiplex FL v2 HRP blocker was applied to each section. The slides were incubated for 15 minutes at 40℃. Slides were washed in PBS/0.2% Triton X-(PBS-T) and blocked with 10% normal donkey serum for 1 hour at room temperature. Primary antibodies were incubated overnight at 4℃ at the indicated dilutions: goat anti-YFP (1:400, Abcam) and rabbit anti-SPC (1:1000, Abcam) in PBS/0.2% Triton X-(PBS-T). After 3x washing, slides were incubated with Alexa Fluor-coupled secondary antibodies for 1 hour at room temperature - goat 488 and rabbit 647 (all Invitrogen, 1:200). After 3x washing, slides were mounted using Prolong Gold with DAPI (Invitrogen). Images used for RNAscope/IF analysis were captured at consistent exposure times, which were determined with negative controls for primary antibodies.

## Figure legends

**Fig. S1.**
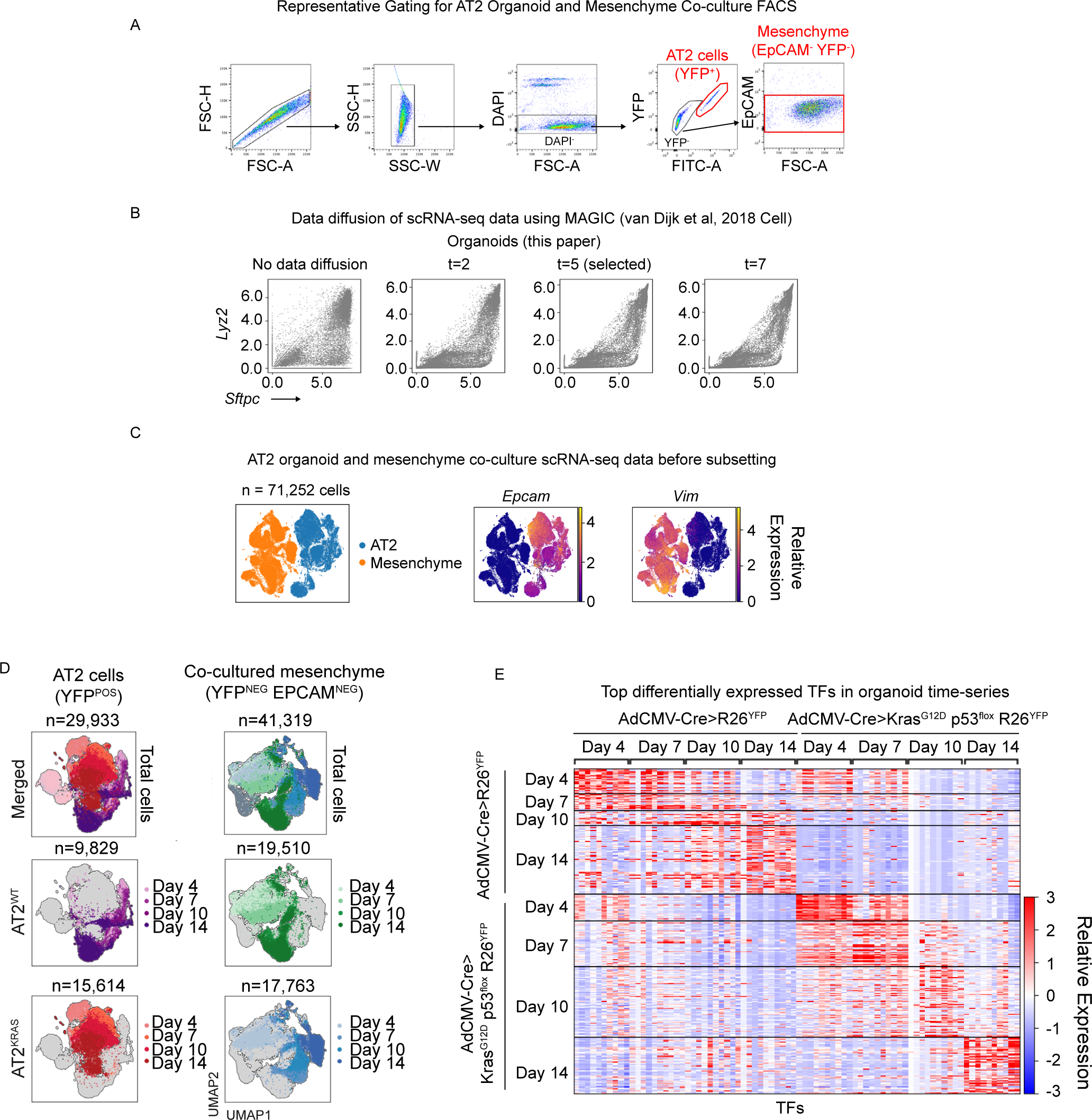
(A) Representative FACS plot for AT2-mesenchyme co-culture time course for scRNA-seq analysis. (B) Correlation between *Sftpc* and *Lyz2* expression in individual organoid co-culture cells, after different levels of data diffusion (t) (*54*). (C) UMAP representation of filtered single cells from organoid co-cultures before subsetting, their corresponding population of origin, and *Epcam* or *Vim* expression. (D) UMAP representations of filtered single cells from organoid co-cultures and their corresponding population of origin, genotype, and time point. (E) Heatmap of the top 10 differentially expressed transcription factors based on genotype and time point.

**Fig. S2.**
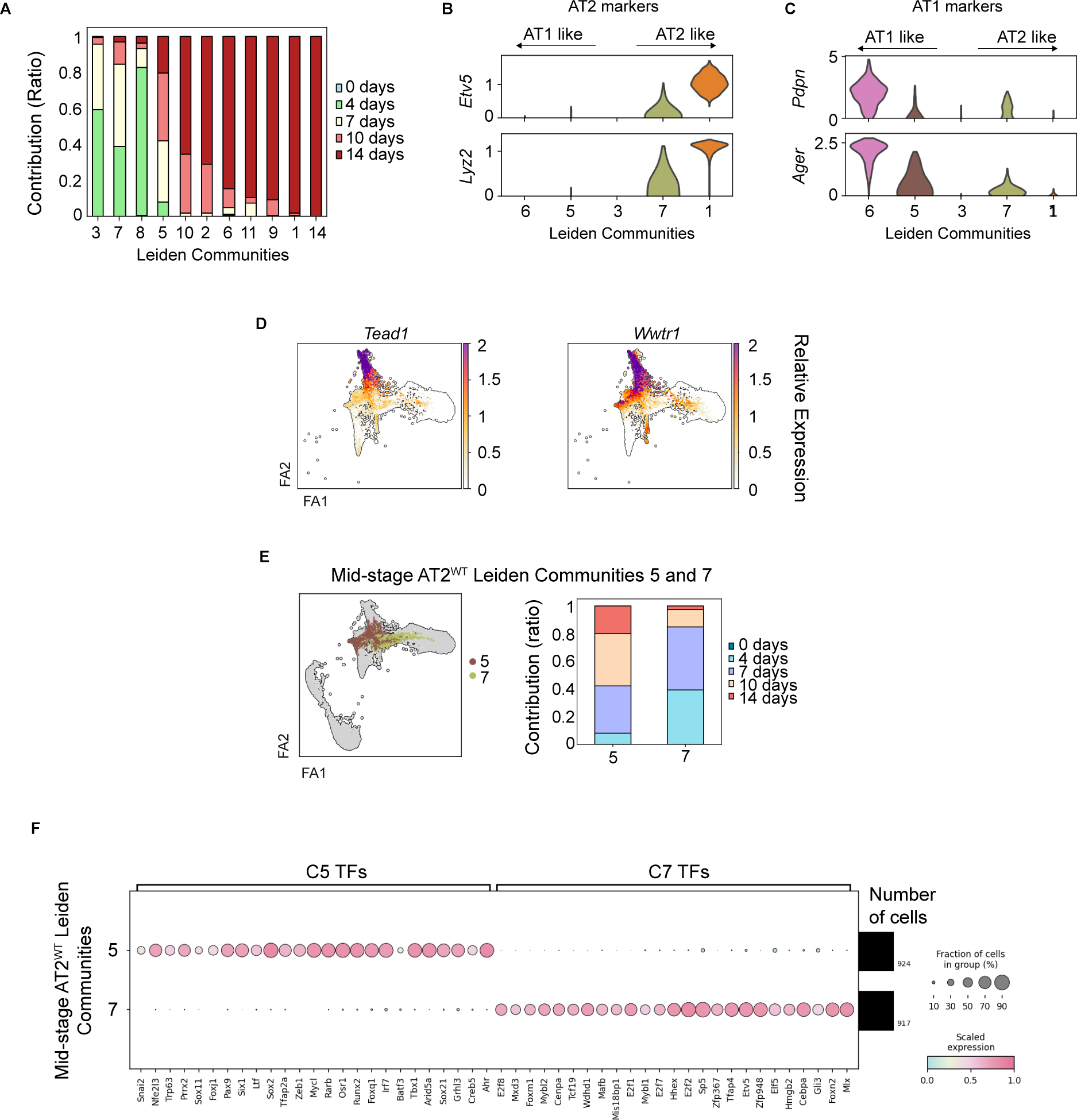
(A) Bar plot representing time point contributions for each Leiden community in the AT2^WT^ organoid scRNA-seq data. (B) Relative expression of AT2 genes *Etv5* and *Lyz2* per community using a violin plot (y-axis, Leiden community; x-axis, relative expression). (C) Relative expression of AT1 genes *Pdpn* and *Ager* per community using a violin plot (y-axis, Leiden community; x-axis, relative expression). (D) FA2 representations of filtered single cells subset from AT2^WT^ organoid data and the relative expression of either *Tead1* or *Wwtr1*. (E) AT2^WT^ organoid intermediates C5 (AT1 fate) and C7 (AT2 fate) within the FA2 space (left). Bar plot representing time point contributions C5 and C7 (right). (F) Relative expression of the top 25 DE TFs in C5 and C7 AT2^WT^ intermediate states using a dot plot (x-axis; DE genes, y-axis; Leiden community).

**Fig. S3.**
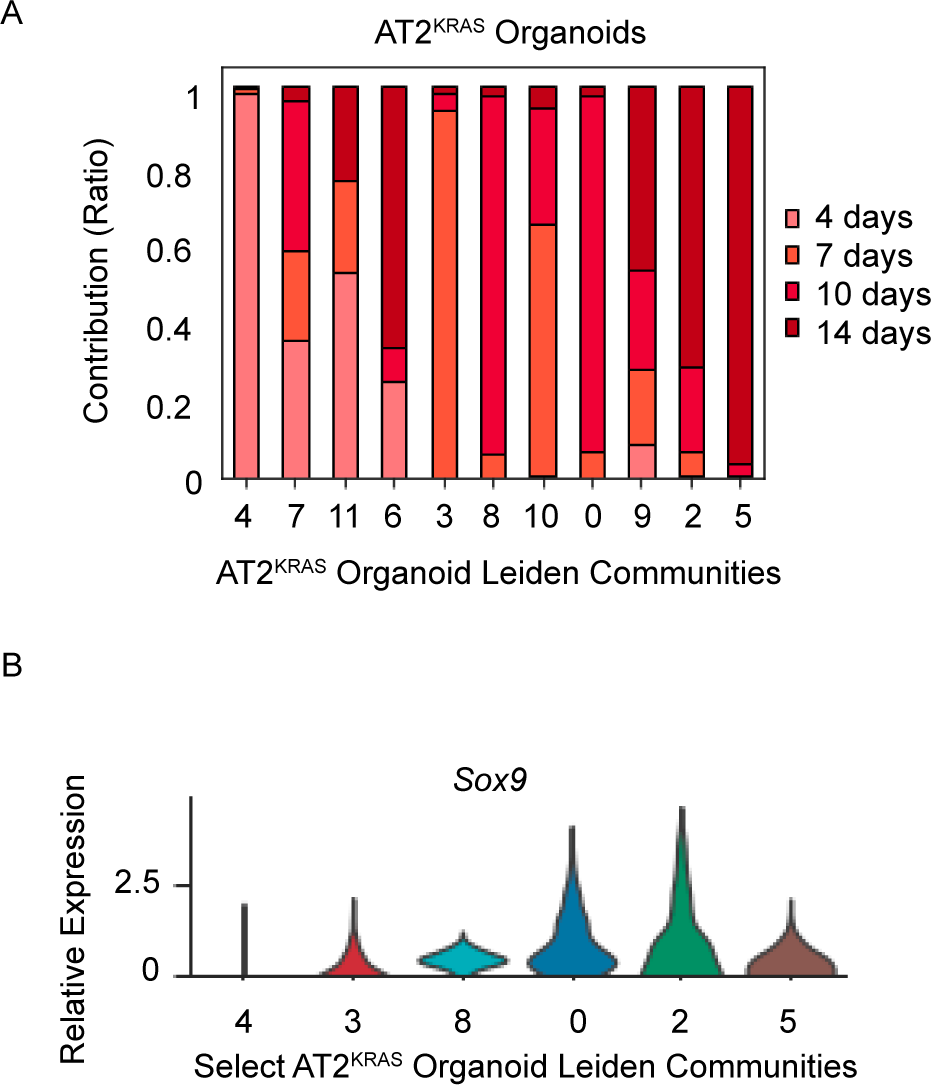
(A) Barplot showing time point contributions in each AT2^KRAS^ organoid Leiden community, represented as a ratio. (B) Relative *Sox9* expression in select early, mid, and late stage communities using a violin plot (y-axis, relative *Sox9* expression; x-axis, Leiden community).

**Fig. S4.**
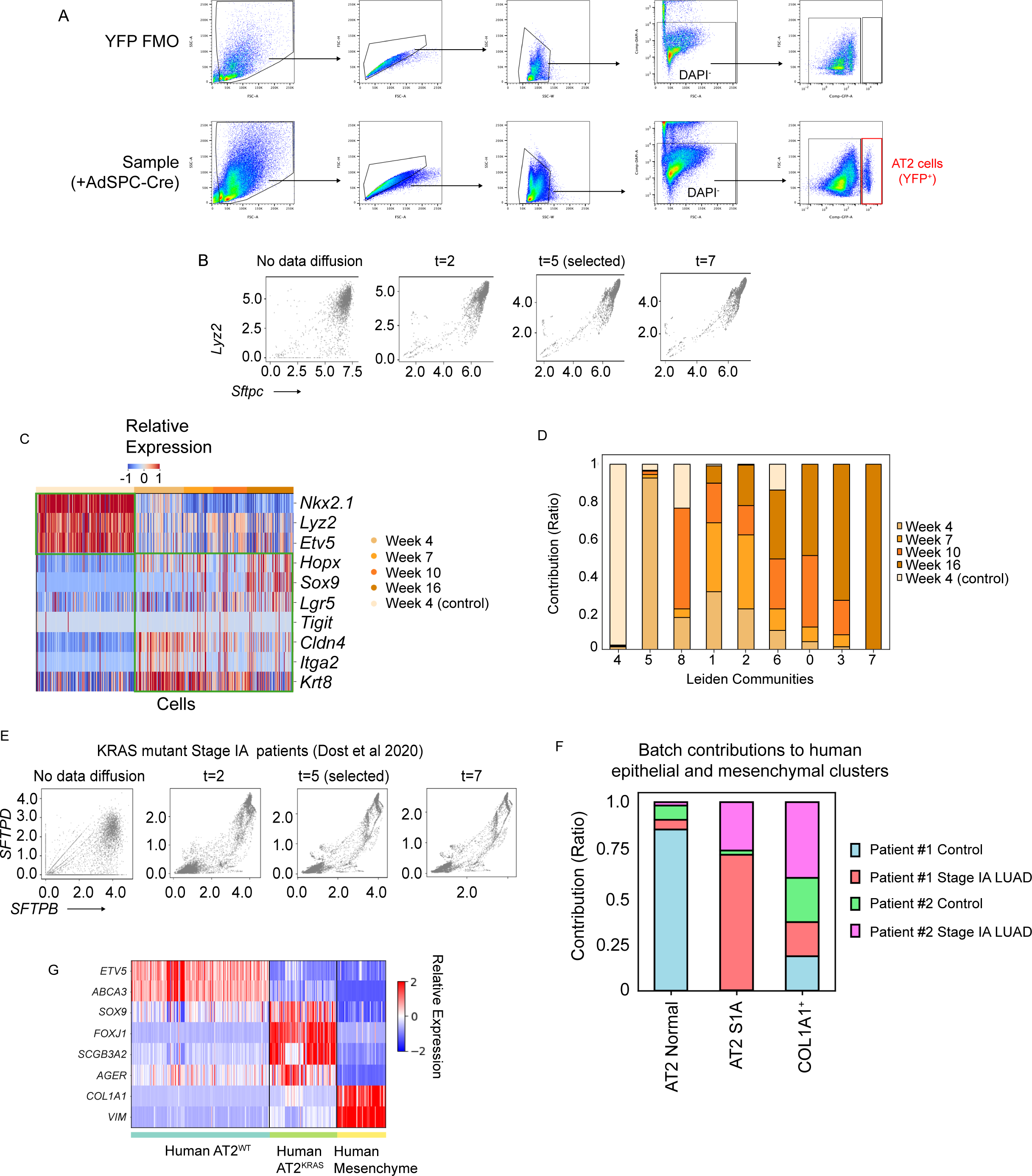
(A) Representative FACS plot for in vivo time course for scRNA-seq experiment. (B) Correlation between *Sftpc* and *Lyz2* expression in vivo, in individual AT2 cells after different levels of data diffusion (t) (*54*). (C) Heatmap of gene expression in in vivo time course data relevant to AT2, AT1, development, stem cell, and injury response gene expression signatures. (D) Bar plot representing time point contributions for each in vivo Leiden community. (E) Correlation between *SFTPD* and *SFTPB* expression in individual stage IA human cells after different levels of data diffusion (t) (*54*). (F) Bar plot representing patient and sample type batch contributions for each Leiden community in the human scRNA-seq data. (G) Heatmap of gene expression in human data relevant to different lung lineages and development.

**Fig. S5.**
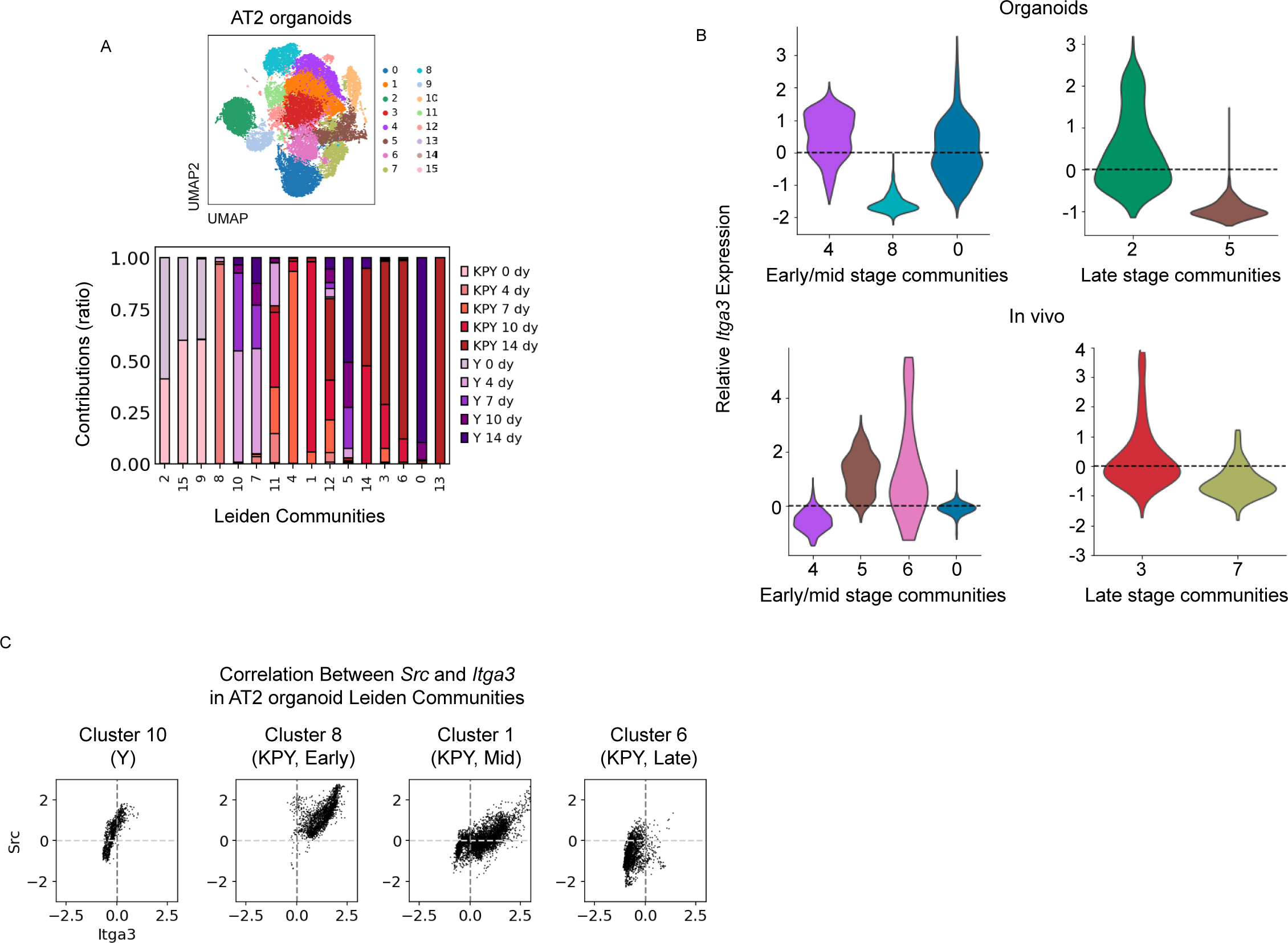
(A) UMAP representation of Leiden communities in the combined AT2 organoid data. Bar plot representing time point contributions for each Leiden community. (B) Relative *Itga3* expression in AT2^KRAS^ organoids (top) and in vivo (bottom) Leiden communities using a violin plot, subset into early/mid and late stage time points (y-axis, relative *Itga3* expression; x-axis, Leiden community). (C) Correlation between *Src* and *Itga3* relative expression at various time points in AT2 organoids, represented as a scatterplot. Each point represents a single cell.

